# Rapid femtosecond laser trenching accelerates cryo-electron tomography lamella workflows

**DOI:** 10.64898/2026.10.23.761368

**Authors:** Christian M. Orr, Pranav N. M. Shah, Éilís C. Bragginton, Saikat Chakraborty, Helen M. E. Duyvesteyn, Stephen Riggs, David I. Stuart, Armin Wagner

## Abstract

Transmission electron microscopy requires electron-transparent specimens, typically below 200 nm thick. For cellular and tissue samples this is typically achieved by focused ion-beam (FIB) thinning to form lamellae. Milling times scale with specimen thickness, and high-pressure-frozen specimens more than 10 µm thick can take up to 2 hours per lamella. We present a pre-processing approach in which bulk material is removed by ultrashort-pulse laser ablation prior to milling, rapidly generating large trenches in vitrified grids under cryogenic conditions and leaving material to be finished by conventional milling to yield electron-transparent lamellae. Laser ablation completes in seconds (4 seconds per pattern), compared with minutes to tens of minutes for FIB trench milling, and introduces negligible damage to the final lamellae. By replacing the most time-intensive, contamination-sensitive bulk-removal steps with seconds-long laser trenching, this open-source approach reduces time-to-lamellae, increasing throughput and preserves compatibility with standard cryo-FIB/SEM polishing and cryo-ET data-collection pipelines.

---

Electron cryo-tomography (cryo-ET) reconstructs three_-_dimensional volumes of biological material in a vitrified, near native state^1^ and, through subtomogram averaging of repeated structures, can reach sub-nanometre detail in situ, as recently demonstrated for cellular ribosomes^2,3^. Cryo-ET is constrained by specimen thickness^4^: direct transmission imaging requires samples to be on the order of a few hundred nanometres, with thicker material suffering from severe multiple scattering, reducing interpretability. Most cellular and tissue specimens therefore require a thinning step to produce electron transparent lamellae suitable for tilt series acquisition.

Typically, samples are thinned with cryo_-_focused ion beam (cryo-FIB) milling^5–7^, in which material is ablated under cryogenic conditions to leave a lamella of 100–200 nm thickness. This has enabled *in-situ* structural cell biology by opening windows into the cell interior while avoiding many of the mechanical artefacts associated with vitreous sectioning^8^. Recent improvements in instrumentation, stability and software control have driven increasing automation of routine milling steps, reducing operator burden and improving reproducibility and throughput^9^.

Despite these advances, conventional plunge_-_frozen, on_-_grid cryo_-_FIB workflows can be compromised by poor vitrification of thicker specimens, low particle/cell density in accessible regions, and low throughput when targets are sparse or small. High pressure freezing^10^ (HPF) is a more effective sample preparation technique for thicker, more concentrated samples as crystalline ice formation is reduced at high pressure. The waffle method^11^ combines HPF with the use of an EM grid as a spacer/support, producing a vitrified slab that can then be milled into large, specimen_-_dense lamellae, providing many of the practical bulk approach advantages while retaining the convenience of on_-_grid preparation and compatibility with manual and automated cryo_-_FIB hardware.

Time-to-lamella is dominated by manual bulk material removal steps, which typically require an operator to remain at the instrument, iteratively adjusting milling parameters and confirming that each site is progressing as intended. Cryo-FIB milling is intrinsically serial and, to preserve specimen vitrification and minimise milling artifacts, ion beam conditions must be conservative, thereby decreasing ablation rates^12^. In waffle workflows this bottleneck is amplified: opening the initial deep trenches and clearing large volumes of vitreous material can be particularly time-consuming, and demands attentive, manual intervention before the automated milling and polishing steps. Recent work also highlights that mechanical failure modes (e.g., cracking/breakage from stress concentration) can materially impact yield, motivating geometry innovations such as stress-relieving features^13^.

To address throughput, the field is progressing from gallium liquid metal FIBs to plasma FIBs (PFIBs), which enable higher beam currents for rapid bulk milling^14^. Xe-PFIB has been demonstrated for bulk milling of high_-_pressure frozen specimens and can support high_-_resolution *in situ* structural biology^9,15^. Systematic analyses have quantified the depth scale over which low beam current PFIB surface damage measurably affects macromolecular information content as less than 50nm^16^. In parallel, commercial instruments have been developed offering automation, high throughput operation, and reduced handling, for example the Thermo Fisher Scientific Arctis Cryo_-_PFIB, with automated sample handling and workflow connectivity. The Arctis is designed to automate lamella production, increase throughput, and simultaneously reduce contamination by enabling more specimens to be loaded (12) and direct transfer of grids to TEM autoloader systems. However, this system is prohibitively expensive for many labs. Moreover, the high beam currents used for rapid bulk milling can themselves impart damage to the underlying vitreous material, so accelerating trench generation on the ion beam is not without quality trade-off.

While PFIBs accelerate bulk removal, the most manual and least automatable step in waffle-style preparation remains the creation of large trenches and clearance volumes. A compelling complementary strategy is ultrashort_-_pulse laser ablation, particularly the femtosecond (fs) regime, which is widely characterised as enabling extremely rapid material removal with strongly reduced heat_-_affected zones compared with longer_-_pulse approaches^17^. In the broader micro_-_machining literature, femtosecond laser ablation achieves milling rates orders of magnitude above ion beam sputtering for large volumes, motivating integrated “LaserFIB” concepts for fast trenching followed by fine FIB polishing^18^, an approach recently extended to resin-embedded biological tissue^19^.

Here, we apply laser ablation directly to waffle_-_prepared cryo-EM grids to create trenches rapidly, reducing the burden from ion-beam milling to final thinning and polishing steps. The setup is relatively affordable, being assembled from off-the-shelf components and controlled by open-source software. Building on the waffle geometry^11^, where large perpendicular trench generation is an explicit, early step, we targeted exactly the portion of the workflow that is most throughput limiting. By shifting bulk removal from ion sputtering to laser ablation, we shortened milling sessions, reduced cumulative contamination exposure, and increased lamella yield per unit time while preserving lamella suitability for downstream cryo-ET and subtomogram analysis.

Using a femtosecond laser at 515 nm wavelength, we reduced trenching times to 4 seconds per lamella site compared with 24 minutes for Ga-FIB and 7 minutes for Xe-PFIB under typical waffle conditions. Laser-trenched grids maintained vitrification and produced cryo-ET-usable lamellae as judged by lamella quality, tilt-series quality, and subtomogram-averaging resolution at 6-9 Å resolution. These results establish laser trenching as a practical throughput multiplier for waffle cryo-ET pipelines, particularly for institutions with limited access to PFIB instrumentation. Single-particle apoferritin control experiments further demonstrate that laser exposure has minimal effect on high-resolution structural information.

## Results

### Femtosecond laser ablation rapidly generates trenches on cryogenic waffle grids

Femtosecond-laser ablation removes material through ultrashort, high-intensity pulses that vaporise the target with negligible thermal diffusion^17^. We built a dedicated platform, the I23 cryogenic laser-shaping system (Figure 1a), in which 515 nm laser beam with 220 fs pulses is steered by a galvanometer scan head onto an EM grid held at cryogenic temperature in a nitrogen cold-gas stream. Grids are automatically transferred by a robotic arm from an 80-sample storage dewar (Methods). The beam size (FWHM ≈ 5.6 µm) defines the smallest feature size, while confined energy deposition yields sharply defined cuts. Trench geometries are defined in software (Figure 1b) and ablated in a single pass. Handling and manipulation of clipped grids relies on standard tools, and grids are clamped using tweezers glued into magnetic bases (Figure S1).

**Figure 1.**
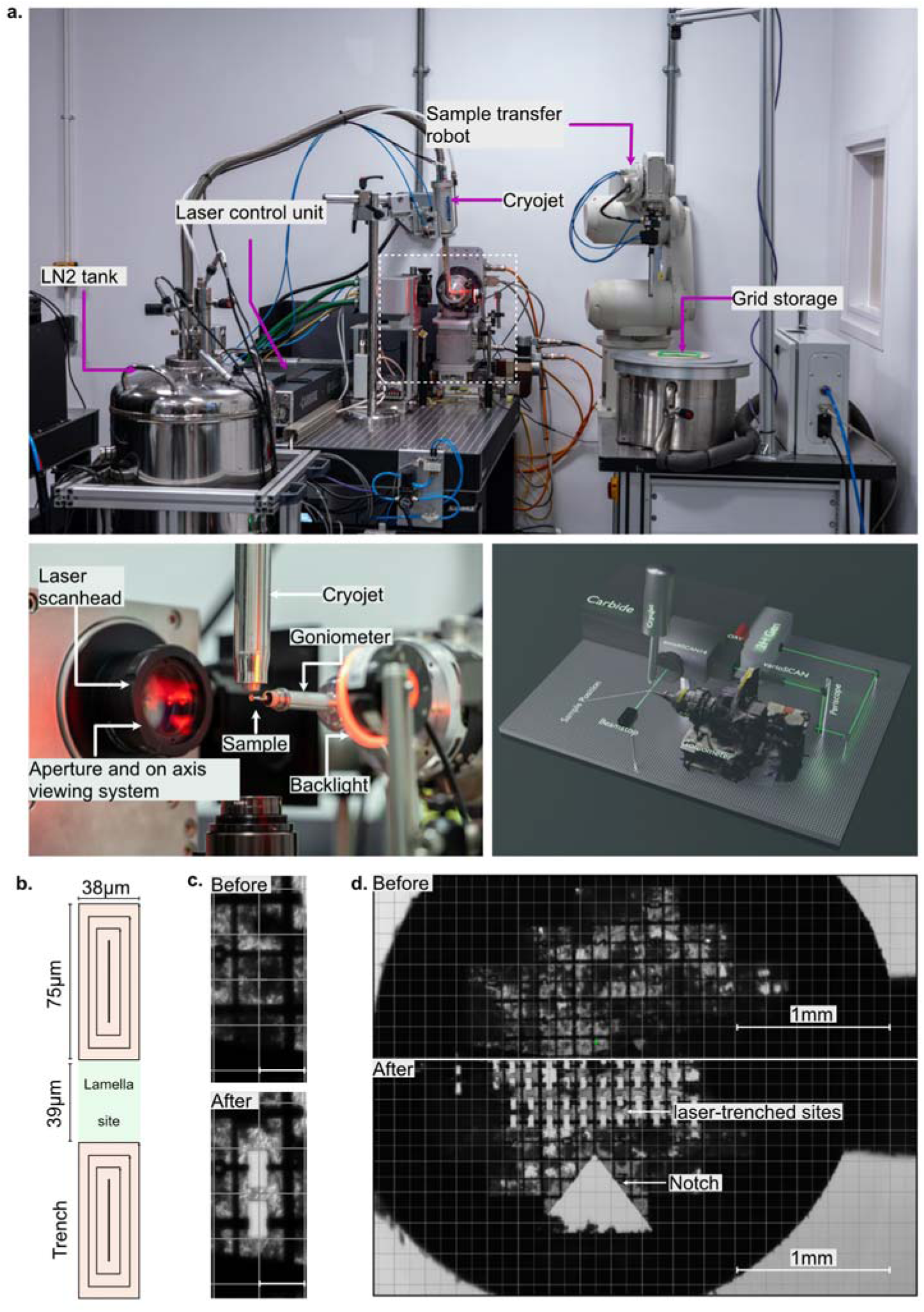
**Femtosecond laser instrument setup and trench generation**. **a.** (top) Overview of the femtosecond laser setup at Diamond Light Source (bottom right) a simplified three-dimensional rendering of the laser-shaping system, showing the principal components on the optical table. (bottom left) enlargement of the inset showing the sample position with respect to the laser, the cryojet and goniometer. **b**. Schematic of the trench milling pattern and its dimensions. **c-d**. Close-up view and overview images of a grid before and after laser milling. Scale bars 100 µm (c.) and 1 mm (d.).

To assess whether femtosecond-laser ablation could replace the bulk-removal step in waffle-grid lamella preparation we generated pre-defined trench geometries directly on high-pressure–frozen cryo-EM grids. Laser trenching reproducibly generated arrays of rectangular trenches across grids (Figure 1c-d).

The trenching geometry was designed to leave a central block of vitreous sample flanked by two rectangular trenches providing adequate clearances and surface for placing lamellae positions on the block at standard FIB milling angles (Figure 1b), we term this the lamella site. To increase efficiency the bottom trench pattern of a row acts as the top trench for subsequent rows of lamella sites. These geometries were defined in the laser-control workflow and executed under cryogenic conditions. To ensure that laser-trenched grids could be loaded into the cryo-FIBSEM instruments in the right orientation, a large triangular notch in a region inaccessible for FIB milling acted as fiducial marker. This cut-out in the grid was visible to the naked eye even when working under liquid nitrogen (Figure 1d).

Inspection of individual sites showed that laser ablation produced cuts without evidence of devitrification, redeposition or microcracking (Figure 1c-d). Patterning was fast across both cell types tested (Gram-positive and Gram-negative bacteria, see Methods). Individual lamella sites were generated in 4 seconds.

A reproducible observation was that front- and back-side surface ice was reduced during laser processing (Figure 1c,d). This surface ice is contaminant frost accumulated during grid handling, HPF and transfer under liquid nitrogen, rather than material introduced by the laser step itself. Ablation is performed at atmospheric pressure within a dry cryogenic gas stream, so ablated debris is carried away from the grid rather than redeposited. Control grids that bypassed the laser step showed no difference in surface contamination, indicating that the additional handling and transfer steps do not introduce measurable ice. For grids contaminated with surface ice, physical and optical access to the underlying vitrified material was improved after generation of the trench geometry (Figure 1c,d).

### Laser trenching increases lamella-site preparation throughput compatible with cryo-FIB and cryo-PFIB polishing

We applied the standard workflow for generating lamellae from grids prepared using the Waffle method^11^ (Figures S2a, S2b). To compare the performance gains using our laser trenching approach, we established the baseline timings for generating lamellae using the Aquilos (Ga-FIB) and Arctis (Xe-PFIB) systems. To facilitate the comparison, all sample preparation conditions and downstream data collection and analyses were identical (Methods).

Semi-automated trench-generation pipelines were developed for the Aquilos and Arctis by defining front and rear rectangular patterns of 22 x 37 µm^2^ and 20 x 17 µm^2^ for the Aquilos and 25 x 40 µm^2^ and 25 x 25 µm^2^ for the Arctis. Trench generation was actively monitored by the operator and stopped once the underlying biological material had been cleared. Dimensions and milling currents were optimised to allow tens of sites to be processed in a 24-hour milling session whilst maintaining beam coherence. Using the Aquilos Ga-FIB, trench generation, bulk clearance and notch-milling took an average of 24 minutes per site whereas the Arctis Xe-PFIB required an average of 7 minutes per site (Figure 2), reflecting the higher sputtering rate of Xe plasma and access to higher coherent milling currents in PFIB systems. Overall, generating a single lamella took approximately 120 minutes on the Ga-FIB system and approximately 53 minutes on the Xe-PFIB system (Figure 2, Table S1). In comparison, laser trenching followed by Ga-FIB polishing reduced the total time to roughly 95 minutes, while laser trenching combined with Xe-PFIB polishing reduced the time to approximately 45 min (Figure 2, Table S1).

**Figure 2.**
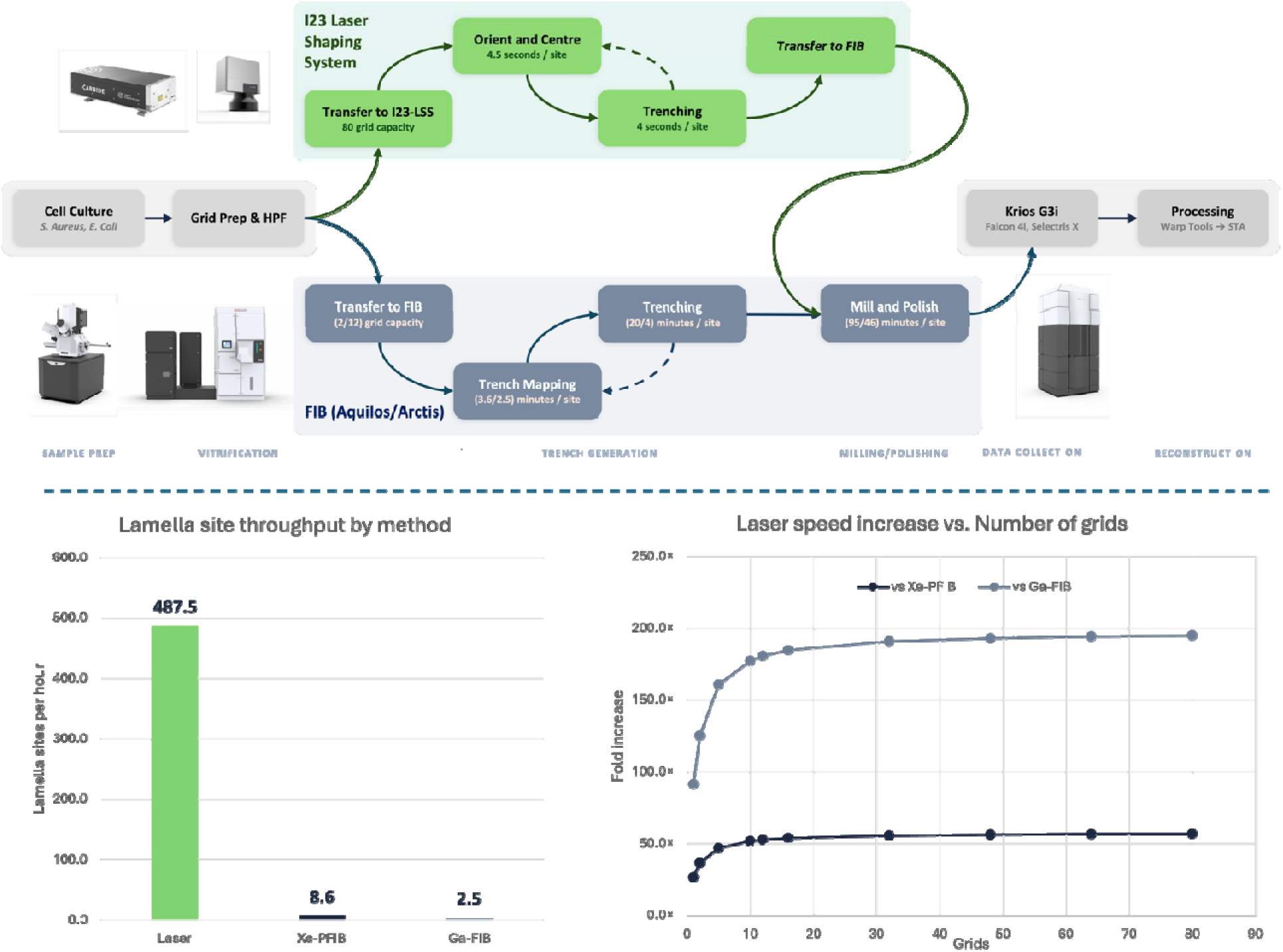
**Hybrid laser–cryo-FIB workflow and per-lamella time comparison. Top**, general workflow for sample preparation. All conditions share common sample preparation (cell culture, grid preparation and high-pressure freezing) and downstream steps. Timings for sample preparations steps are included. **Bottom left**, bar chart comparing the number of lamella sites produced in 1 hour using the I23 Laser Shaping System, the Arctis Xe-PFIB and the Aquilos Ga-FIB. **Bottom right**, speed fold increase using the I23 Laser Shaping System compared to Aquilos Ga-FIB and Arctis Xe-PFIB.

Considering the total session time, we developed a timing model for the laser trenching workflow to analyse throughput changes with increasing batch size. In our setup, the laser-shaping dewar accommodates five pucks, each containing 16 grids, corresponding to a maximum capacity of 80 grids. Each grid yielded approximately 80 lamella sites. The model incorporates both the laser trenching time and the operational overheads associated with robot transfers, cooldown and alignment. The model uses 6.2 seconds per lamella site, which incorporates both centring and lamella generation by trenching sites above and below, and assumes 80 lamella sites being cut per grid. Laser trenching was estimated to be 195-fold faster than Ga-FIB trenching on the Aquilos and 57-fold faster than Xe-PFIB trenching on the Arctis assuming a full 80 grid session of just over 13 hours. Substantial gains were also achieved at smaller batch sizes. Processing of only two Classification: Internal grids produced a 125-fold improvement over the Aquilos Ga-FIB workflow in 31 minutes, while a 12-grid session of just over 2 hours, representing a single load for the Arctis, achieved a 53-fold improvement over Xe-PFIB trenching. These results demonstrate that the throughput advantage of laser trenching increases with batch size but remains considerable even for relatively modest numbers of grids.

Grids with laser-generated lamella sites were transferred into Aquilos and Arctis instruments. Milling, thinning and polishing yielded electron-transparent lamellae comparable in appearance and handling to those produced by conventional workflows (Figure 3a-c).

**Figure 3.**
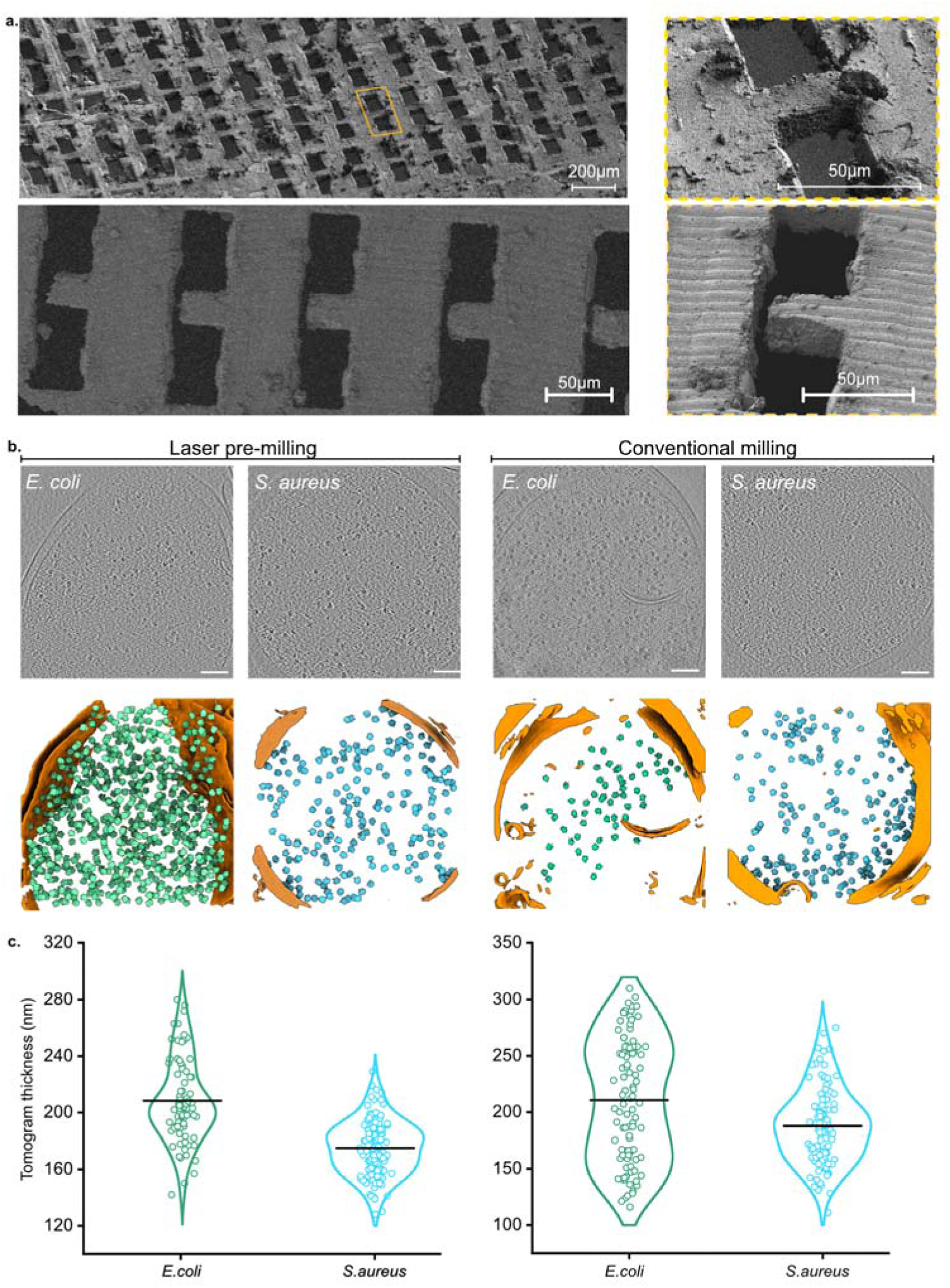
Cryo-FIB milling of a laser-trenched site to electron transparency. **a**. Ion tile-set overviews (Upper row: Arctis, lower row: Aquilos) of a laser-trenched grid showing an array of lamella sites; yellow box enlargement of a candidate site selected for subsequent milling steps. **b.** 5 nm thick virtual section through tomograms (top) and corresponding segmentations (below) identifying ribosome (green; E. coli and blue; S. aureus) positions and bacterial cell membranes (orange). **c.** Violin plots of tomogram thicknesses across different datasets and conditions. Black line indicates mean value.

Across the sample types examined, we observed no increase in lamella loss or mechanical failure. These data indicate that laser trenching can be integrated upstream of either Ga-FIB or Xe-PFIB finishing steps offering a substantial speed up in lamella generation.

### Laser trenching preserves tomographic and subtomogram-averaging quality

To assess whether laser trenching affects specimen quality, we first evaluated the struc ural integrity of reconstructed tomograms. Using the same processing workflow (Methods), comparable cellular morphology, membrane contrast and macromolecular density were observed in both laser-trenched and conventionally prepared lamellae. No reconstruction artefacts could be attributed to laser treatment (Figure 3b).

Laser-trenched lamellae prepared from *Staphylococcus* samples yielded 95 tomograms with a mean thickness of 175 ± 19 nm, *Escherichia coli* lamellae yielded 100 tomograms with a mean thickness of 208 ± 29 nm (Figure 3c). Conventionally prepared lamellae from *Staphylococcus* samples yielded 128 tomograms with a mean thickness of 187 ± 32 nm, and *Escherichia coli* lamellae yielded 138 tomograms with a mean thickness of 213 ± 56 nm. The resulting tilt series supported high-quality three-dimensional reconstructions of both Gram-positive and Gram-negative bacteria, in which the distinct cell-envelope architectures and intracellular macromolecular organization of the two species were readily resolved (Figure 3).

Abundant 70S ribosomes were identifiable within the cytoplasm of both species, enabling us to assess the preservation of molecular structural information by subtomogram averaging (Figure 3). 70S ribosome reconstructions were obtained from laser-trenched and conventionally prepared lamellae. Following particle picking, classification and particle refinement, high-quality ribosome reconstructions were obtained from all four datasets, with global resolutions of 6.9 Å and 6.3 Å for the laser-trenched *E. coli* and *Staphylococcus* cells respectively and 9.32 Å and 8.58 Å for the corresponding conventionally prepared lamellae (Figure 4, Table S2). In every reconstruction, the large ribosomal subunit was resolved to higher local resolution (close to Nyquist limit of ∼3.8 Å) than the small subunit, consistent with the greater conformational variability of the latter. Overall map quality, global resolution and local-resolution distributions were comparable.

**Figure 4:**
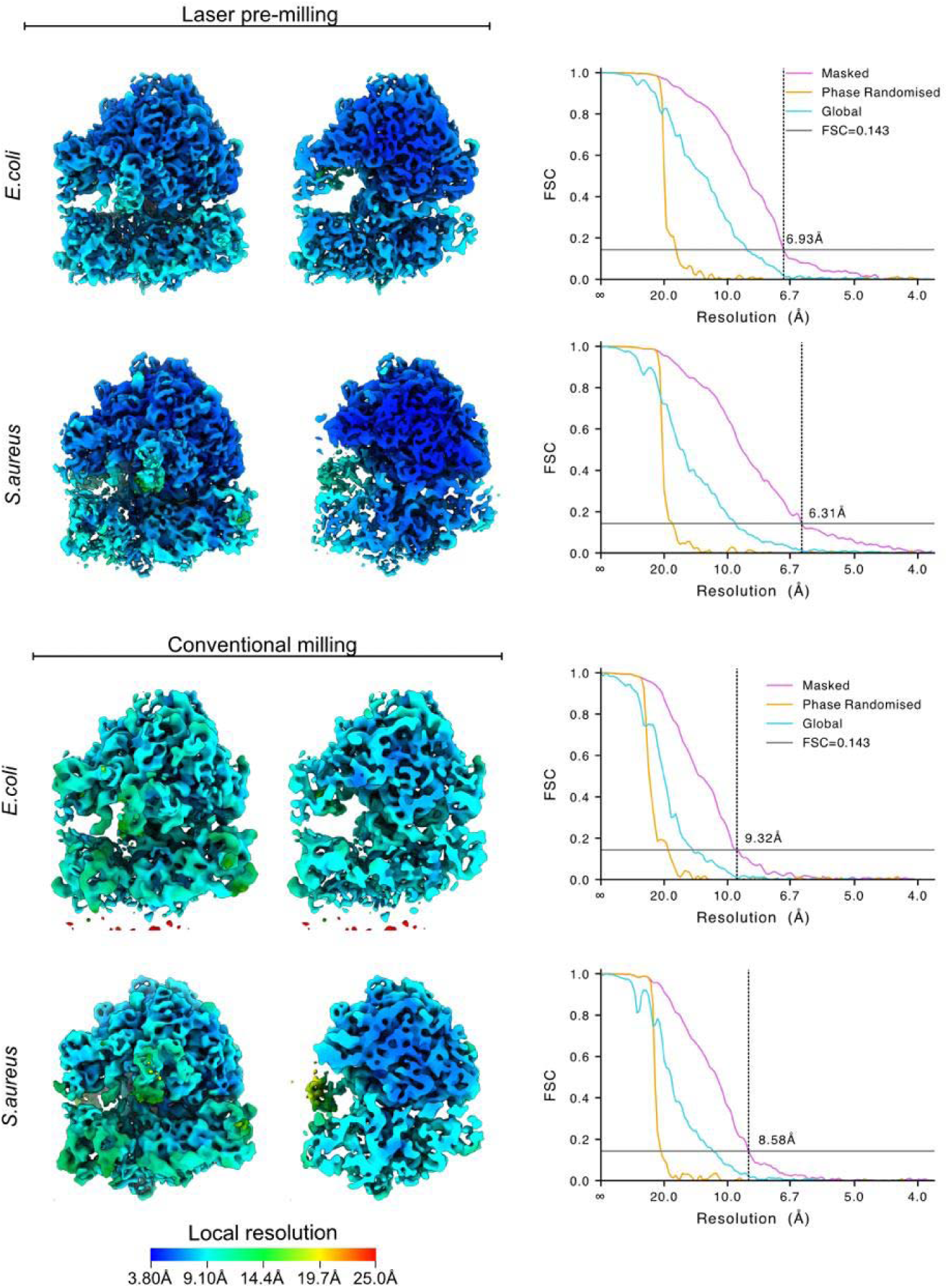
Sub-tomogram averaging of ribosomes from laser-trenched and conventionally prepared samples. Ribosome isosurfaces are coloured using local resolution values (see scale bar) and global Fourier shell correlation (FSC) curves are depicted. Resolutions are reported at the gold-standard FSC cutoff of 0.143.

Together, these results indicate that replacing ion-beam trenching with laser trenching does not produce a detectable loss of tomographic reconstruction quality or subtomogram-averaging performance. Laser trenching therefore preserves biologically relevant structural information while increasing the throughput of lamella preparation.

### Femtosecond laser trenching minimally affects high-resolution structural information

To evaluate potential laser damage on high resolution structural information, we exposed single-particle cryo-EM grids containing human apoferritin (ApoF) to the laser under conditions mimicking those above. Grid squares were cut using either rectangular or circular patterns (Figure 5a, Table S3). Grids that were not exposed were processed in parallel and served as controls.

**Figure 5.**
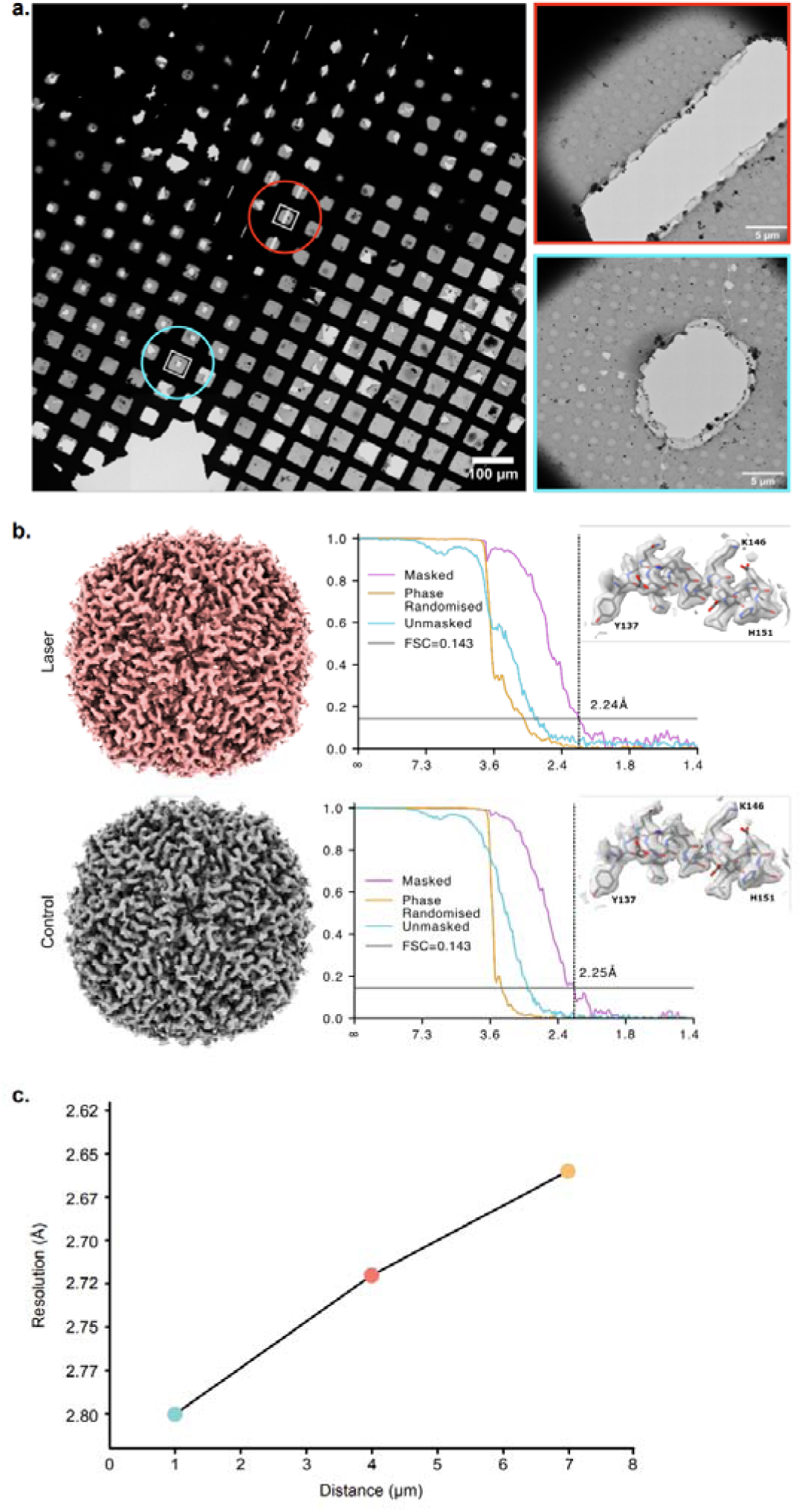
Effect of laser exposure on high resolution structural detail. a, Low magnification overview of a vitrified grid with laser milled line (red) and circular patterns (cyan) **b**, Isosurface representations (contoured at _σ_=3) of human apoferritin. Coulomb potential maps together with the Fourier shell correlation plots and a closeup region of a helix showing residues 137 – 151 with PDB: 7R5O docked by rigid-body fitting in UCSF ChimeraX^20^. **C**, Plot showing the change in apoferritin resolution as a function of particle distance from the laser-ablated area.

Following standard single-particle data collection and processing (Methods), the laser exposed dataset yielded a consensus ApoF reconstruction at 2.24 Å resolution from a final curated set of 76,689 particles while an equal number of particles from the unexposed control dataset reached a comparable resolution of 2.25 Å (Figure 5b, Table S3). Laser milling did not, overall, produce a detectable loss of high-resolution structural information.

We next assessed whether reconstruction resolution varied as a function of particle distance from the edge of the laser-ablated area. Particles were divided into distance-based subsets of 2 µm, and an independent reconstruction was generated for each subset of the same particle number (n=10 822). Particles within the first 0 – 2 µm of the laser-ablated area reconstructed to a global resolution of 2.8 Å with a resolution loss of 0.16 Å in comparison to the reconstruction of the non-lasered control (2.64 Å). The global resolution improved for particles 3 – 5 µm from the laser-ablated area to 2.72 Å and within 6 – 8 µm the resolution is comparable to the control with a global resolution of 2.66 Å (Figure 5c). Overall, these results show laser-ablation to cause minimal damage.

## Discussion

Femtosecond laser trenching removes a major bottleneck from waffle-style lamella preparation for cryo-ET. In our workflow, initial trenching is accelerated up to 195-fold, with 80 lamella sites per grid produced in under 10 minutes. After laser processing, grids are transferred into a FIB instrument, where the operator selects sites of interest and uses automated thinning and polishing workflows to produce the final lamellae. Because conventional trench generation requires several minutes of supervised ion-beam time, replacing this step with the laser substantially increases throughput (Figure 3a). The laser ablates copper and gold as readily as vitreous material, allowing trenches to cross grid bars, giving access to shallow tilt angles that are otherwise restricted in waffle workflows (Figure 1d). The trenches generated here were more than twice the length and 5-fold and 3.5-fold the area of conventionally milled grids on the Aquilos and Arctis, respectively.

High-current FIB trenching can compromise material surrounding the milled region. In particular, high-current xenon plasma FIB milling has been shown to generate an amorphous layer at the milled surface. Even with successively reduced current polishing of lamella, substantial structural damage extends some 30 nm into the specimen, becoming negligible from around 45 nm^16^.

Conventional high-current trenching can also redeposit sputtered material and devitrify the surrounding sample, limiting the density at which lamella sites can be prepared. By contrast, laser trenching is performed at atmospheric pressure in a cold gas stream, avoiding redeposition of ablated material onto the grid, and we observed no evidence that laser ablation caused sample devitrification. This permits lamella sites to be positioned at higher density and makes more efficient use of the specimen, allowing more lamellae to be prepared from a single grid.

No significant laser-induced damage was observed. Visual inspection of the lamella sites revealed no devitrification, redeposition or cracking, and laser-trenched specimens yielded tomograms and ribosome reconstructions comparable to conventionally prepared controls. Single-particle analysis of apoferritin similarly showed little effect on overall resolution within a few micrometres of the edge of a laser-ablated hole. The modest reduction in resolution observed closest to the ablated region may reflect reduced mechanical support and increased specimen movement acting in combination with radiation damage during imaging. Importantly, the final lamella is thinned from the central block and is therefore spatially separated from the laser-machined trench walls, excluding potentially affected material from the imaged volume. Any residual laser-induced damage must also be considered in the context of damage introduced during final ion-beam polishing which itself produces a structurally damaged surface layer extending tens of nanometres into the completed lamella^16,21,22^. Thus, the absence of a detectable loss of structural information in laser-trenched lamellae indicates that laser ablation does not introduce damage that limits subsequent high-resolution cryo-ET.

The system was originally designed to shape protein crystals for the long-wavelength macromolecular crystallography beamline I23 at Diamond Light Source^23^. Assembled from off-the-shelf components and driven by open-source control software, such a system could be readily made available at large imaging centres. Our approach to machining cryogenic specimens is not restricted to the rectangular geometry used here. Pillars, cylinders and stress-relieving features that would be prohibitively slow to generate by ion milling become readily accessible, as does the shaping of vitrified specimens for ptychography, soft X-ray imaging and crystallography. More generally, laser ablation is immediately applicable to cryogenic specimen-preparation workflows in which removal of large volumes of material is a major bottleneck, including lift-out approaches for preparing lamellae from bulk specimens. We regard this workflow as a starting point and anticipate that hybrid laser-FIB preparation will become a routine route to high-throughput sample production for cryo-ET and other techniques where large volumes of materials need to be removed with minimal damage.

## Data availability

Coulomb potential maps have been deposited at the Electron Microscopy Data BANK (EMDB) under accession codes: EMD-59459 (laser-milled *Staphylococcus aureus* 70S ribosome), EMD-59460 (laser-milled *Escherichia coli* 70S ribosome), EMD-59464 (Conventionally milled *Staphylococcus aureus* 70S ribosome), EMD-59470 (Conventionally milled *Escherichia coli* 70S ribosome), EMD-59473 (laser-exposed apoferritin), EMD-59474 (apoferritin control).

Scripts to perform distance-based partitioning of apoferritin particles can be accessed from https://github.com/stephen-riggs/laser-map-segment

## Methods

### Cell culture

A glycerol stock of TOP10 *Escherichia coli* cells was inoculated into 10 ml LB broth and grown overnight at 37 °C. Cells were harvested by centrifugation at 4,000 × *g* for 5 min. *Staphylococcus aureus* cells were grown from a glycerol stock by inoculation into 10 ml YPD medium, cultured for 48 h at 30 °C and harvested under the same conditions.

### Grid preparation and high-pressure freezing

Two high-pressure freezing protocols were used depending on instrument availability.

For vitrification with the HPM-100 (Leica Microsystems), aluminium 3 mm type B planchettes (Leica Microsystems) were coated with 2% (w/v) soya lecithin in chloroform and left to evaporate shortly before use. Harvested cells were washed in 1 mL PBS, pelleted by centrifugation (4,000 xg, 2 min), resuspended in 1 mL PBS containing 10% (v/v) glycerol, pelleted again under the same conditions and resuspended in the remaining supernatant at half the pellet volume. Copper H2 finder grids with a lacey carbon film (Agar Scientific) were glow discharged in air for 30 seconds using an easiGlow (Ted Pella). Five microlitres of cell suspension were applied to each grid immediately before high-pressure freezing.

For vitrification with the HPM-010 (Bal-Tec), aluminium planchettes (3 mm, 0.3/0.50 mm; Wohlwend) were sanded with fine-grit paper until no visible surface defects remained, washed with 70% ethanol, coated with 2% (w/v) soya lecithin in chloroform and left to dry for 1-2 hours. Shortly before freezing, cells were centrifuged gently (300g) for 2 minutes and resuspended in half their volume. Gold 200-mesh grids with 2/1 spacing and a carbon support film (Agar Scientific) were plasma cleaned in air for 1 minute (Harrick Plasma). Five microlitres of cell suspension were applied to each grid before high-pressure freezing.

### Femtosecond laser shaping system

The laser source was a CARBIDE CB5 6W ytterbium laser (Light Conversion), air-cooled, with a fundamental wavelength of 1,030 ± 10 nm. A harmonics generator attached to the output provided access to the second (515 nm) and third (343 nm) harmonics. In this study the second harmonic was used exclusively, avoiding potential infrared-regime heating and ultraviolet-regime absorption effects. At 515 nm the maximum output power was 3 W, with a maximum pulse energy of 50 µJ, a minimum pulse duration of approximately 220 fs and a fundamental repetition rate of 60 kHz divisible by 1–1,000 (minimum 60 Hz).

The beam was directed via four mirrors to a water-cooled excelliSCAN 14 galvanometer steering system (SCANLAB) equipped with a varioSCAN de II 20i dynamic focus unit before the main scan-head beam-input port and an f = 65 mm f-theta objective lens with 515-nm-compatible coating. The galvanometer mirrors were water-cooled. An on-axis viewing system was installed using a SCANLAB 515-nm-compatible camera adapter and an Alvium G1-1240M-CH-C camera (Allied Vision).

### Grid handling and transfer

High-pressure-frozen grids were transferred under liquid nitrogen to custom cryogenic sample holders using standard vitrified-grid handling tools and a clipping station. Each holder comprised a SPINE pin base^24^ fitted with negative-action tweezers (WPI, catalogue no. 501779-G) secured with Stycast adhesive, providing stable cryo-compatible support throughout manipulation. Mounted grids were loaded into CombiPucks for transfer into the laser-shaping robot dewar. Inside the dewar, a BART robotic system^25^ transported each holder to the goniometer stage, where a CryoJet^26^ maintained cryogenic temperature during laser-shaping operations. When necessary, minor adjustments to grid orientation were performed manually using cryo-cooled tweezers. Following laser processing, grids were returned to grid boxes via the BART robot by reversing the transfer sequence.

### Laser shaping parameters

Sample visualisation, alignment and image capture were performed using Aithre^27^, a custom PyQt-based graphical interface. Laser operating parameters were set via the CARBIDE User App using the second harmonic (515 nm) at 21% transmission and a 10-kHz repetition rate. A crosshair displayed on the on-axis viewer defined the laser focal position, and the target grid square was positioned accordingly. Beam steering and execution of cutting routines were controlled using LaserDESK (SCANLAB), in which trench geometries were imported from predefined .dxf or .las pattern files. The mirror translation speed was set to 0.02 m s⁻¹, and each pattern was cut in a single pass, typically requiring 4 seconds depending on geometry. After each pass, the grid was re-evaluated to assess cut completeness; additional passes were performed when required. Once the desired trench profile was achieved, the next grid square was selected and the procedure repeated. To facilitate rapid identification of grid orientation during subsequent loading into cryo-FIB/SEM instruments, a large triangular fiducial aperture was additionally ablated through the grid as an unambiguous orientation marker.

### FIB milling – Aquilos

Clipped grids, with or without prior laser shaping, were loaded into an Aquilos 2 (Thermo Fisher Scientific) using a 35° pre-tilt shuttle. Grids were screened using the microscope user interface, and surface ice contamination was removed by FIB imaging at 7 nA and 30 kV with the grid perpendicular to the ion beam. Grids were subsequently sputter-coated with metallic platinum for 60 s (1 V, 30 mA) followed by a 50 second GIS platinum deposition and a second sputter coat under the same conditions.

For laser-shaped grids, low-magnification montages were collected in MAPS v3.22, and lamella positions were selected after visual inspection. Sites were imported into AutoTEM v2.4 and milled in an automated manner at a milling angle of 12–16 °. Lamella sites were placed at the leading edge with milling patterns extended above and below to remove bulk material. Rough milling was carried out at 3.0 nA and the lamella was further thinned stepwise to a target thickness of 200 nm in AutoTEM, with the current progressive reduced to 0.1 nA. Final polishing was performed in two steps at 50 pA and 30 pA, with a target thickness of 110 nm in AutoTEM.

For lamella prepared by conventional waffle milling, trenches were generated as previously described^11^. Placing the grid perpendicular to the ion beam, two trenches were milled at the top (22 x 37 µm) and bottom (20 x 17 µm) with a spacing of 25 µm in the centre of the grid square at 16 nA to generate the site for lamella generation. After trench milling, bulk material was manually cleared from the bottom of the lamella site at an initial milling angle of 60 ° using a current of 7 nA, followed by a milling angle of 40 ° and 20 ° using a current of 3 nA. After manual clearance of the lamella sites, a notch was inserted to the right of the lamella at 1 nA^28^. Lamella sites were manually added to AutoTEM and final lamella generated in an automated fashion as described above.

### FIB milling – Arctis

Clipped grids were loaded into an Arctis cryo-plasma FIB (Thermo Fisher Scientific). Using the Web UI 1.6^29^ interface, grids were screened automatically by collecting grid-overview images in SEM mode. Ice contamination on the front side of each non-lasered control grid was removed by Xe-PFIB imaging at 15 nA and 30 kV, held at a shallow angle (approximately 12–14°) and at the lowest magnification available, until most of the ice had been cleared. The front side of each laser-trenched and control grid was then coated with ion-sputtered metallic platinum for 120 seconds (12 kV, 70 nA), followed by a 100-s GIS platinum deposition and a second 120 seconds sputter coat under the same settings.

For laser-trenched grids, lamellae-placement sites were visually inspected in ion tile-set views to identify smooth clean top surfaces, free of contamination and were then marked for lamellae generation in the Web UI. Each site was milled in an automated manner using xenon at 30 kV and a milling angle of 15°. Lamella were placed on the top surface at the leading edge (Figure S2a), while milling patterns were extended to cover material top and bottom of the lamella surface. Rough milling was carried out at 1.0 nA. The lamella checkpoint in Web UI 1.6 was used to repeat the rough-milling step when remnant material underneath the lamella was not fully cleared. However, the large clearance volume generated by the laser design allowed material to ablated in one go in most cases. After rough milling, the resulting 2.5 µm blocks were thinned down stepwise with progressively reduced milling current. Final polishing was performed in two steps, at 30 pA followed by 10 pA. A target thickness of 150 nm yielded final lamellae of approximately 200–250 nm after polishing (Figure 3c).

For conventional waffle milling, two trenches were milled at each lamella site from the front side of the grid, with the stage tilted to 10°, corresponding to a FIB incidence angle of 48° relative to the grid plane. The front and rear trenches measured 40 and 25 µm in height, respectively, with both trenches measuring 25 µm in width. Trenches were milled using a 60 nA Xe plasma beam at 30 kV, maintaining a separation of approximately 20–25 µm between the two trenches at each site. Following trench milling, lamella locations were identified in WEB UI 1.6 using SEM and/or ion tile-set images acquired at eucentric height. Subsequent lamella milling was performed using the same workflow and milling conditions described above for the laser-trenched grids. The final lamella thickness ranged from 200 to 250 nm, as determined by manual measurements in IMOD v5.1.

### Cryo-ET data acquisition

Tilt series were acquired on a Krios G3i microscope equipped with a Selectris X energy filter and a Falcon 4i direct electron detector (Thermo Fisher Scientific). Data were collected using a dose-symmetric tilt scheme^30^ with the PaceTomo^31^ script in SerialEM^32^. A tilt range of ±54° was used with 3° increments, beginning at ±15° depending the upon orientation of lamella front after loading to account for the lamella pre-tilt angle. Target defocus was varied across a range of −3 to −5 µm in 0.5-µm steps. Images were recorded in EER mode on the Falcon 4i at a nominal pixel size of 1.9026 Å and a total dose of approximately 3.0 e⁻/Å² per tilt. A 10-eV slit width was applied throughout acquisition. Tracking and focusing were performed before and after each tilt image. The zero-loss peak of the energy filter was tuned once before each data-collection session.

### Tomogram reconstruction, visualisation and subtomogram analysis

Data were processed using WarpTools^33^. EER files were grouped at approximately 1 e⁻/Å² per frame for motion correction and simultaneous CTF estimation. Motion was modelled with a 1 × 1 × 5 spatial–temporal grid owing to the low per-frame dose. The CTF was estimated using a window size of 768 pixels and fitted to a resolution of 4 Å, with a 2 × 2 × 1 grid to account for the defocus gradient across tilted projections. Even and odd frame averages were saved for subsequent denoising.

Tilt series were imported into WarpTools ^33^ (dev39) with a tilt-axis angle of 84.7° and a pre-tilt offset of 15° subtracted from all tilt angles. Tilt stacks were generated at a pixel size of 10 Å by Fourier cropping. Tilt series alignment was performed with AreTomo v.1.3.4^32,34–36^ using a minimum field-of-view criterion of 0.8 followed by MissAlignment^37^ using developer shared parameters. Alignment parameters were imported into WarpTools at the corresponding pixel size^32–36^. Defocus handedness was determined automatically over 10 tilt series and corrected prior to per-tilt CTF refinement.

CTF-corrected tomograms were reconstructed at a pixel size of 10 Å for particle picking. For denoising, even and odd tomograms were reconstructed and processed using WarpTools^33^. Tilt-series were visually scored for their interpretability and amount of ribosome they contained. Badly aligned or thick tomograms were discarded from further processing resulting in a final dataset as listed in the Table S2. Tomogram thickness was measure manually in IMOD^38^ by measuring the distance between the top and bottom surfaces along different views and averaged. Membrane segmentations were performed using Membrain-Seg using default parameters and visualised in ChimeraX^20^. Particle picking was performed using TomoCPT^39^ with pre-trained general weights and a particle diameter of 280 Å on contrast-inverted tomograms. Particles were extracted as three-dimensional subtomograms at a pixel size of 5 Å and a box size of 96 pixels using WarpTools (Table S2, Figure S3). Initial poses were determined by 3D classification in RELION v4.0.2^40^2 into 5 classes with a regularisation parameter of T = 4. Classes lacking structural detail were discarded, and the retained particles (see Table S2) were subjected to 3D auto-refinement in RELION with a spherical mask of 300 Å diameter.

Refined particles were imported into M^2^. A population, source and species were defined against the original tilt-series settings, and the reference map was resampled to 1.9026 Å per pixel. Two rounds of multi-particle refinement were performed: the first refining particle poses and the second additionally refining image-warp with a 3 × 3 image-warp grid. CTF defocus parameters were refined once the reconstruction reached approximately 8 Å resolution. Thus, refinement of the electron-optical parameters simultaneously with tilt-geometry optimisation yielded ribosome reconstruction with global resolutions of 6.3 Å, 6.9 Å, 8.58 Å, 9.32 Å respectively (Table S2, Figure 4).

### Apoferritin Grid Preparation

Human heavy-chain apoferritin was purified as previously described^41^.

Purified apoferritin was vitrified by plunge freezing 3 µl of 2 mg ml^-1^, applied to a glow discharged R1.2/1.3 300 mesh copper grid (Quantifoil) and blotted for 3 s. The grids were frozen using a Vitrobot Mark IV (Thermo Fisher Scientific) operating at 4 °C, 100 % Humidity, blot force 3.

An apoferritin grid was subsequently transferred, and a combination of single points or lines were shaped into the grid. The transfer, grid handling and laser shaping parameters used are identical to those described above for the High-Pressure Frozen samples. A grid which did not undergo any additional handling or laser shaping was kept as a control.

### Apoferritin Data Collection

For the laser-shaped apoferritin grid, data was collected on either a Titan Krios G3 (Thermo Fisher Scientific) at the Electron Bioimaging Centre at Diamond Light Source (eBIC) or on a Titan Krios G3i (Thermo Fisher Scientific) at the Oxford Particle Imaging Centre. Both microscopes were fitted with a Falcon 4i Direct Electron Detector and a Selectris X energy filter (Thermo Fisher Scientific). Due to issues with automated hole recognition, data was collected using Tomo5 v5.21 (Thermo Fisher Scientific) using targeted single acquisition. A defocus range of –0.5 to –3.0 µm was targeted in 0.3 µm increments. Data were recorded as TIFF files at a nominal pixel size of 0.723 Å, with a total dose of 50 e⁻/Å². A 10 eV slit width was applied throughout data acquisition.

For the apoferritin control grid, data was collected on a Titan Krios G3 (Thermo Fisher Scientific) at eBIC, Diamond Light Source, fitted with a Falcon 4i Direct Electron Detector and a Selectris X energy filter (Thermo Fisher Scientific). Data was collected using EPU v3.11 (Thermo Fisher Scientific) using the same data collection parameters as above.

### Apoferritin Data processing

Single-particle datasets from the different data collections were imported into RELION 5.0^40,42,43^, with each dataset assigned to a separate optics group. Raw movies were grouped into virtual frames corresponding to a cumulative dose of 1 e /Å² per frame, gain-corrected, and subjected to RELION’s implementation of MotionCor2 using a 5 × 5 spatial grid, followed by CTF parameter estimation using CTFFIND-4.1^44^.

Individual apoferritin particles were identified using an in-house-trained crYOLO^45^ general particle-picking model. Extracted particles were initially down sampled to a pixel size of 5.84 Å/px and subjected to 2D classification. Particles from the selected classes were then re-extracted at the physical pixel size in 512-pixel boxes and underwent a round of 3D classification with octahedral symmetry imposed. Classes displaying clear secondary-structure features were subjected to gold-standard FSC-based 3D refinement with octahedral symmetry imposed.

Electron-optical parameters were subsequently refined iteratively, including beam tilt, per-particle defocus and for the control dataset, anisotropic magnification and 4^th^ order aberrations. Finally, reference-based particle motion correction was performed, followed by a final 3D refinement, yielding consensus reconstructions at resolutions of 2.24 Å for the laser-exposed dataset and 2.25 Å for the control dataset (Table S3).

### Distance-based particle partitioning of Apoferritin Data

To measure the specific contribution of laser exposure to the amount of recoverable particle resolution, a custom Python script was developed to assign particles according to their distance from the laser-ablated area. For each search map, the laser-ablated area was identified using MemBrain**-**seg^46^. Search-map images were padded to a common size and down sampled prior to segmentation, and the resulting segmentation was rescaled to the original search-map dimensions. Connected regions within the segmentation were identified and the largest connected component was retained as the mask representing the laser-exposed region.

Acquisition positions associated with each search map were extracted from the microscope BatchPositionsList.xml metadata. Their stage coordinates were transformed into search-map pixel coordinates using the stage, image-shift, and reference transformation matrices stored in the corresponding SearchMap.xml file. This transformation also accounted for the search-map pixel size, coordinate orientation, and the position of the map centre. Particle acquisition positions were then associated with their corresponding search maps and transformed using these corrections.

For each acquisition position, the Euclidean distance to every pixel belonging to the segmented laser-exposed region was calculated, and the minimum value was taken as the distance from that position to the nearest boundary of the exposed region. Pixel distances were converted to physical distances using the search-map pixel size and acquisition binning. These distances were subsequently appended to the corresponding particle metadata (rlnHelicalID), allowing particles to be partitioned according to their distance from the laser-ablated area for downstream reconstruction and resolution analysis.

Using the calculated distance information, particles were partitioned into distance-based bins according to their proximity to the laser-exposed region. For each distance bin, half-maps were reconstructed independently using the RELION command-line program *relion_reconstruct*. The resulting half-maps were subsequently processed using *relion_postprocess* to estimate the average reconstruction resolution and the B-factor used for map sharpening.

## Acknowledgements

We thank Cecilia Rocchi of Grimes lab for providing *Escherichia coli* culture, Weixian Lu (Strubi) and James Birch (Membrane Protein Laboratory, Diamond Light Source) for providing cells, and Karl Harlos from Strubi for assistance with transport logistics. We also thank Cecilia Pozzi and Marco Mazzorana for the apoferritin and Gabriel Karras (Diamond Light Source) for help with laser alignment.

Electron microscopy experiments and sample preparation were performed at both the Oxford Particle Imaging Centre, OPIC, an Instruct centre, and the UK national electron Bioimaging Centre (eBIC) at Diamond Light Source, proposal CM44121-1. Computational aspects of this work were enabled by the Oxford Biomedical research Computing (BMRC) facility.

## Author contributions

D.I.S., A.W. and C.M.O. conceived and developed the idea of using the laser shaping system as part of the cryo-ET workflow. É.C.B, S.C, H.M.E.D, and P.N.M.S performed the cell culture, high pressure freezing and grid preparation. C.M.O and A.W conceptualized and built the laser shaping system. C.M.O developed the grid handing and SPINE mount adapters and performed the laser shaping experiments. É.C.B and S.C. performed cryo-FIB experiments and collected cryo-ET data. É.C.B and S.C analysed and visualised cryo-ET data. É.C.B prepared the apoferritin grids, collected the data and É.C.B and P.N.M.S carried out apoferritin SPA. S.R wrote the scripts for the distance-based partitioning of the apoferritin data. The initial draft was written by C.M.O, review and editing of the paper were carried out by all authors. Funding for the laser shaping system was acquired by A.W.

