## Supplementary Information for "Rapid femtosecond laser trenching accelerates cryo-electron tomography lamella workflows"

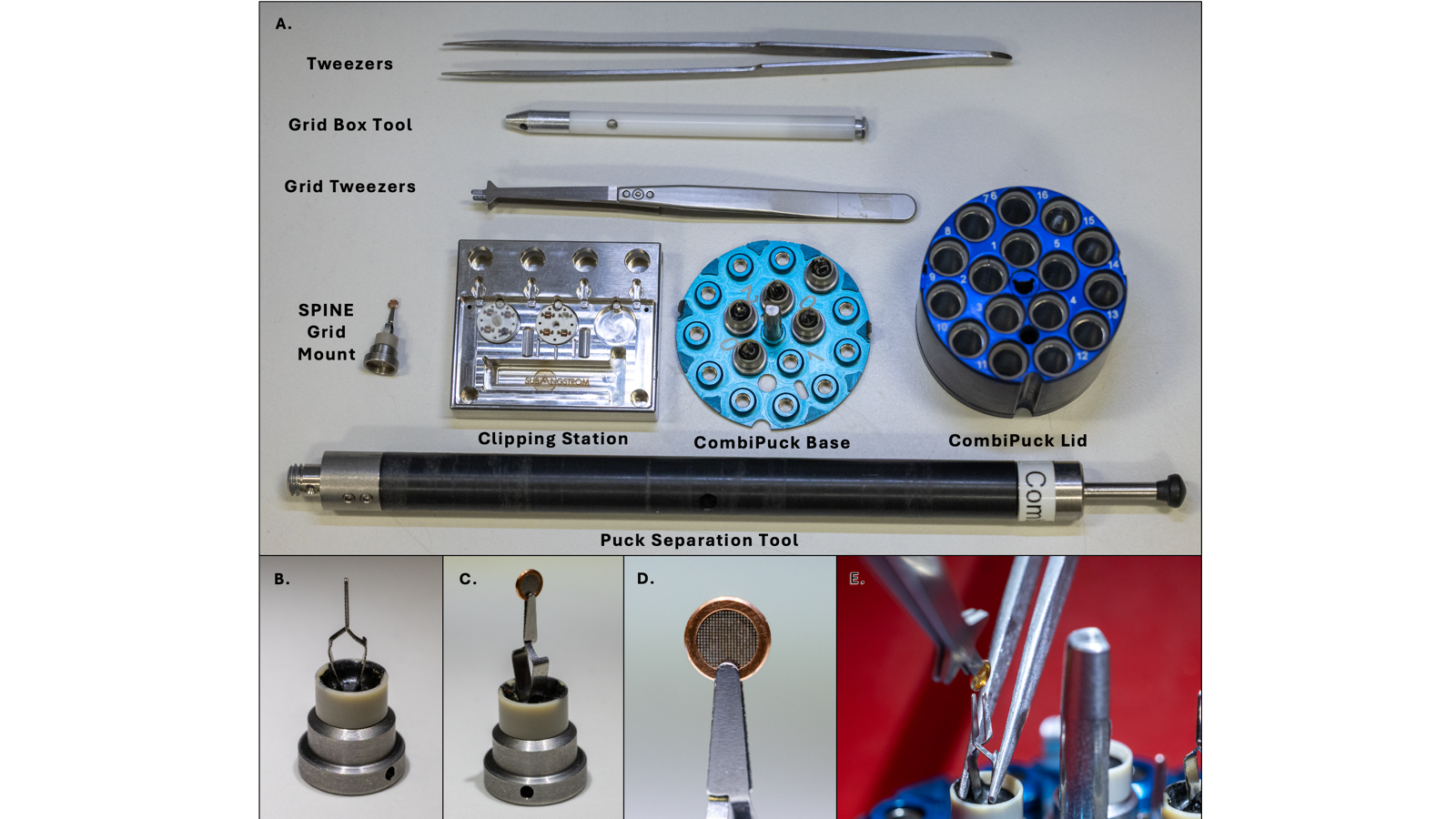
***Figure S1*** **Instrumentation for grid handling. A.** Tweezers and grid manipulation and storage hardware. **B.-D.** Close-up of the spine mounted clipped grids. **E.** Using T-tweezers to place clipped grids in the modified spine.

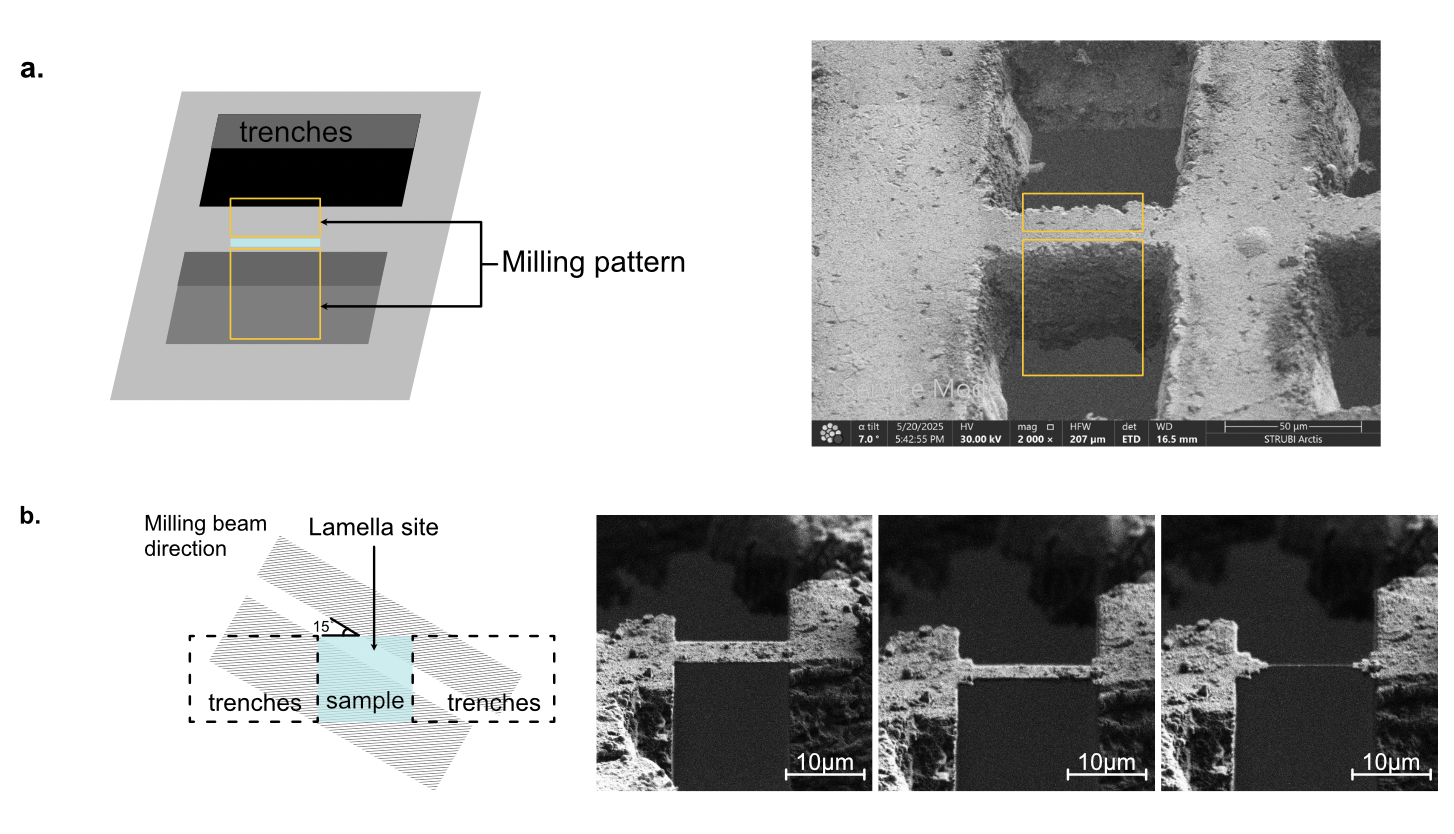
***Figure S2***. **Lamellae generation from Waffle grids. A.** Schematic (left) and SEM image (right) of trenches generated using either the laser or the conventional methods and placed pattern for subsequent milling. **B.** Milling pattern and subsequent step wise thinning of the material.

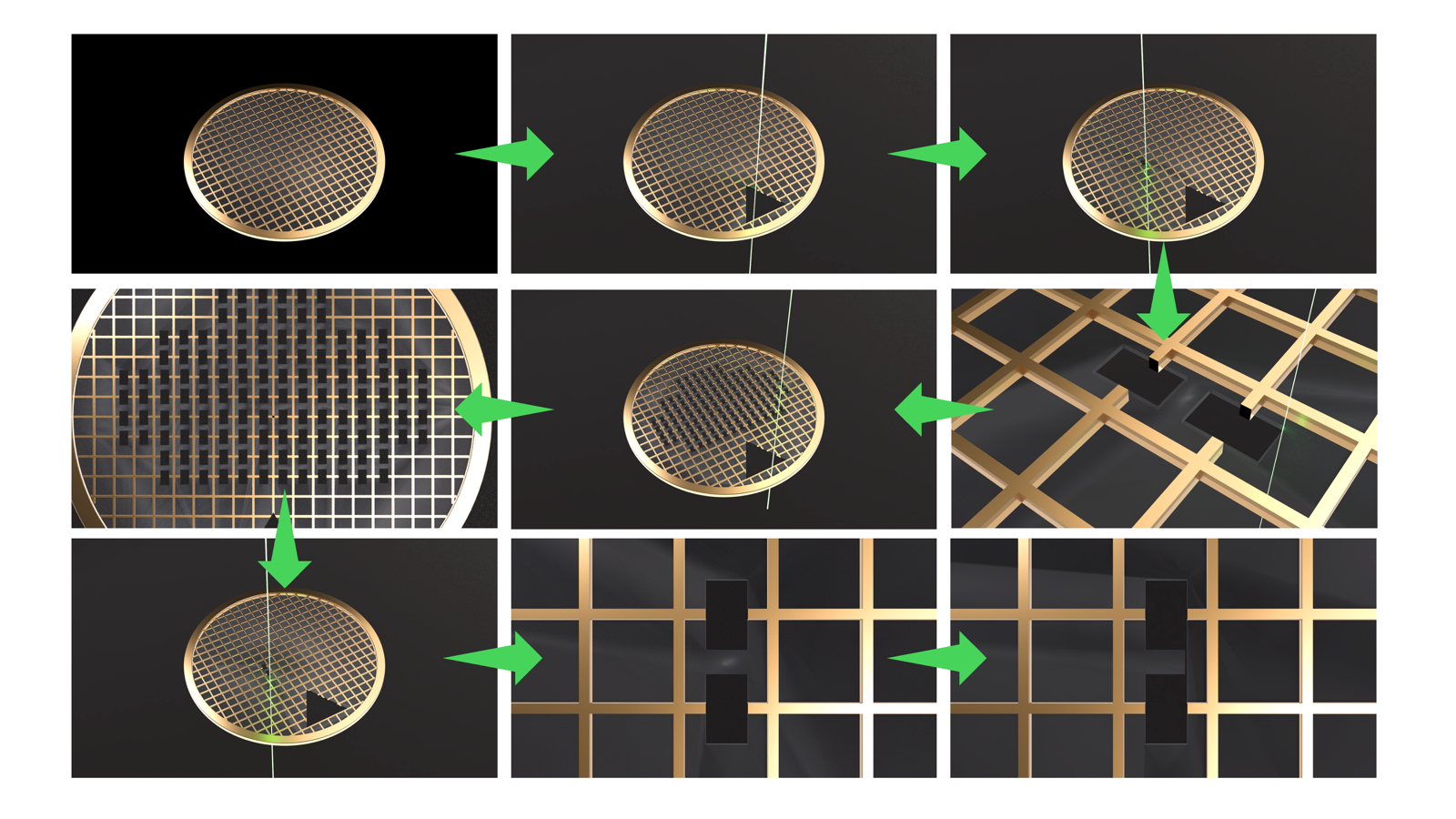

***Figure S3.*** **Illustration of laser ablation workflow.**

***Table S1.*** Laser pre-trenching throughput as a function of batch size.

| Pucks | Grids | Lamellae | Laser (min) | Xe-PFIB (min) | Ga-FIB (min) | Speed-up vs Xe-PFIB | Speed-up vs Ga-FIB |
| --- | --- | --- | --- | --- | --- | --- | --- |
|  | 1 | 80 | 21 | 560 | 1,920 | 26.7× | 91.4× |
|  | 2 | 160 | 31 | 1,120 | 3,840 | 36.5× | 125.2× |
|  | 5 | 400 | 60 | 2,800 | 9,600 | 46.9× | 160.9× |
|  | 10 | 800 | 108 | 5,600 | 19,200 | 51.7× | 177.2× |
|  | 12 | 960 | 128 | 6,720 | 23,040 | 52.6× | 180.5× |
| 1 | 16 | 1,280 | 166 | 8,960 | 30,720 | 53.9× | 184.7× |
| 2 | 32 | 2,560 | 322 | 17,920 | 61,440 | 55.7× | 191.0× |
| 3 | 48 | 3,840 | 477 | 26,880 | 92,160 | 56.4× | 193.2× |
| 4 | 64 | 5,120 | 632 | 35,840 | 122,880 | 56.7× | 194.3× |
| 5 | 80 | 6,400 | 788 | 44,800 | 153,600 | 56.9× | 195.0× |

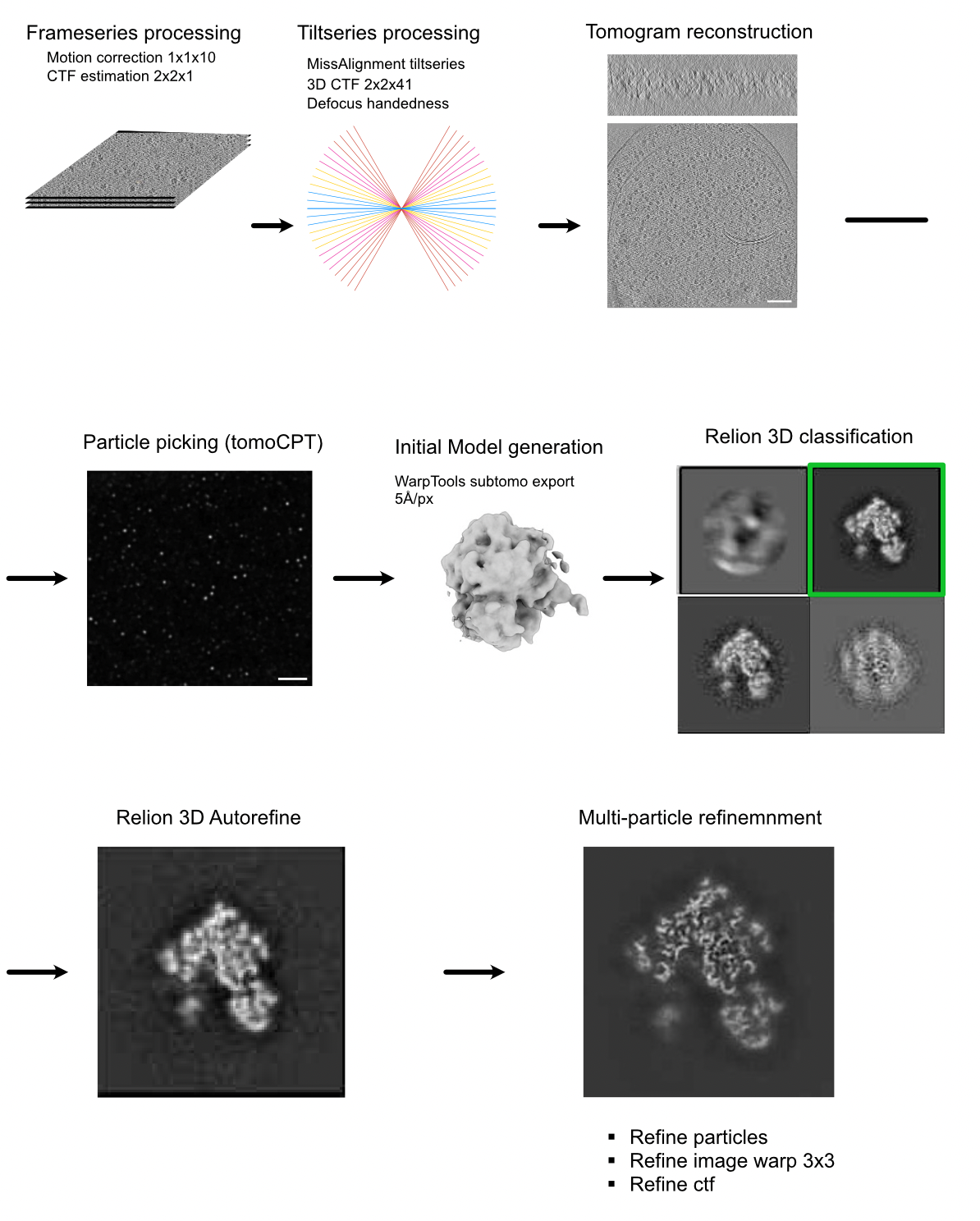

***Figure S4.*** **Workflow for obtaining subtomogram averaging structure of 70S ribosome.**

***Table S2.*** Statistics for cryo-ET data collection and processing

| Sample | *E. coli* | *Staphylococcus* | *E. coli* | *Staphylococcus* |
| --- | --- | --- | --- | --- |
| *Milling* | | | | |
| Laser ablation | Yes | | No | |
| Gas for milling | Xe | Xe | Xe | Xe |
| Milling angle (°) | 15 | 15 | 15 | 15 |
| *Data collection* | | | | |
| Voltage (kV) | 300 | 300 | 300 | 300 |
| Detector | Falcon IV | Falcon IV | Falcon IV | Falcon IV |
| Energy filter | Selectris X | Selectris X | Selectris X | Selectris X |
| Data acquisition software | SerialEM-PaceTomo | SerialEM-PaceTomo | SerialEM-PaceTomo | SerialEM-PaceTomo |
| Pixel size (Å/px) | 1.9026 | 1.9026 | 1.9026 | 1.9026 |
| Total electron dose across tilt series (e^-^/Å^2^) | 118 | 108 | 118 | 112 |
| Dose (e^-^/Å^2^) | 3 | 3 | 3 | 3 |
| Tilt range (°) | -51 to 60 | -51 to 54 | -51 to 60 | -51 to 57 |
| Frame number | 182 | 287 | 301 | 196 |
| Acquisition scheme | Dose symmetric | Dose symmetric | Dose symmetric | Dose symmetric |
| Defocus range (µm) | -3 to -5 | -3 to -5 | -3 to -5 | -3 to -5 |
| Number of tilt-series | 100 | 95 | 138 | 128 |
| *Processing* | | | | |
| Processing software | WarpTools | WarpTools | WarpTools | WarpTools |
| Number of tomograms | 100 | 95 | 138 | 128 |
| Tomograms used for STA | 56 | 50 | 78 | 68 |
| Number of initial subtomograms | 35422 | 18441 | 42482 | 23900 |
| Number of subtomograms after classification | 35421 | 6224 | 4013 | 7416 |
| Symmetry imposed | C1 | C1 | C1 | C1 |
| Au-standard resolution at 0.143 FSC (Å) | 6.91 | 6.32 | 9.3 | 8.58 |
| EMDB code | EMD-59460 | EMD-59459 | EMD-59470 | EMD-59464 |

***Table S3.*** Statistics for apoferritin data collection and processing

| **Sample** | **Apoferritin – laser ablated** | **Apoferritin – control** |
| --- | --- | --- |
| ***Data collection*** | | |
| Voltage (kV) | 300 | 300 |
| Detector | Falcon IV | Falcon IV |
| Energy filter | Selectris X | Selectris X |
| Data acquisition software | Tomo5 | EPU |
| Pixel size (Å/px) | 0.723 | 0.723 |
| Total dose (e^-^/Å^2^) | 50 | 50 |
| Defocus range (µm) | -0.5 to -3.0 | -0.5 to -3.0 |
| ***Processing*** | | |
| Processing software | Relion 5.0 | Relion 5.0 |
| Number of micrographs | 6048 | 5111 |
| Initial particle images | 254, 131 | 187, 823 |
| Final particle images | 76,689 | 76,689 |
| Symmetry imposed | O | O |
| Au-standard resolution at 0.143 FSC (Å) | 2.24 | 2.25 |
| EMDB code | EMD-59473 | EMD-59474 |
